# Towards a mesoscale physical modeling framework for stereotactic-EEG recordings

**DOI:** 10.1101/2022.07.06.498826

**Authors:** Borja Mercadal, Edmundo Lopez-Sola, Adrià Galan-Gadea, Mariam Al Harrach, Roser Sanchez-Todo, Ricardo Salvador, Fabrice Bartolomei, Fabrice Wendling, Giulio Ruffini

## Abstract

**Objective:** Stereotactic-EEG (SEEG) and scalp EEG recordings can be modeled using mesoscale neural mass population models (NMM). However, the relationship between those mathematical models and the physics of the measurements is unclear. In addition, it is challenging to represent SEEG data by combining NMMs and volume conductor models due to the intermediate spatial scale represented by these measurements.

**Approach:** We provide a framework combining the multicompartmental modeling formalism and a detailed geometrical model to simulate the transmembrane currents that appear in layer 3, 5 and 6 pyramidal cells due to a synaptic input. With this approach, it is possible to realistically simulate the current source density (CSD) depth profile inside a cortical patch due to inputs localized into a single cortical layer and the induced voltage measured by two SEEG contacts using a volume conductor model. Based on this approach, we built a framework to connect the activity of a NMM with a volume conductor model and we simulated an example of SEEG signal as a proof of concept.

**Main results:** CSD depends strongly on the distribution of the synaptic inputs onto the different cortical layers and the equivalent current dipole strengths display substantial differences (of up to a factor of four in magnitude in our example). Thus, the inputs coming from different neural populations do not contribute equally to the electrophysiological recordings. A direct consequence of this is that the raw output of neural mass models is not a good proxy for electrical recordings. We also show that the simplest CSD model that can accurately reproduce SEEG measurements can be constructed from discrete monopolar sources (one per cortical layer).

**Significance:** Our results highlight the importance of including a physical model in NMMs to represent measurements. We provide a framework connecting microscale neuron models with the neural mass formalism and with physical models of the measurement process that can improve the accuracy of predicted electrophysiological recordings.

## 1. Introduction

Stereotactic-electroencephalography (SEEG) is a technique in which intracerebral implanted electrodes are used to record brain electrical activity in the form of electric potentials. This technique is routinely used in patients with drug-resistant epilepsy in order to determine the spatiotemporal organization of the epileptogenic network within the brain and identify potential targets for surgical resection [1].

There is a wide agreement that extracellular potentials which can be measured by electroencephalographic (EEG) methods or other techniques such as local field potentials (LFPs), in the low frequency range (0 to a few kHz), are to a large extent created by the transmembrane currents induced by synaptic inputs (post-synaptic currents) onto pyramidal cells. This is due to the coherence in space (alignement “in palisades” of pyramidal cells) and time [2] of these currents, which allows them to contribute in an additive manner into the extracellular voltage, as well as to the elongated form factor of pyramidal cells, which lead to large current dipoles.

When neurotransmitters bind to synaptic receptors, ion channels on the membrane open and membrane currents flow in or out of the cell to hyperpolarize or depolarize the postsynaptic neuron. An excitatory synapse produces an inward current that is seen as a negative current source (i.e., a sink) from the extracellular medium. Conversely, an inhibitory synapse causes an outward current that is seen as a positive current source (i.e., a source) from the outside. Within the timescale of SEEG recordings, these current sinks or sources are always balanced by a passive return current to achieve electroneutrality. The synaptic currents and their associated return currents can give rise to sizable dipole currents (or higher order n-poles) and, if many such currents synchronize with a coherent direction, a measurable voltage perturbation is produced in the volume conductor.

In the past decades, the activity measured by SEEG electrodes has been studied using mesoscale brain models such as neural mass models (NMMs) [3, 4]. These models are mathematical simplified representations of the dynamics of interconnected neuronal populations [5, 6, 7]. They describe the average neural population dynamics (firing rates, membrane potential perturbations) by capturing relevant physiological features at the mesoscale. To connect with electrophysiology measurements, the average membrane potential is often used as a surrogate of LFPs [8], SEEG [4] or EEG measurements (see, e.g., [9]). More recently, a laminar NMM framework using a simple physical model has been used to represent electrophysiological measurements such as LFPs [10, 11] or SEEG [12]. Here we start from this work to provide a rigorous framework combining computational brain models with the microscale physics of current generation in realistic neuron compartment models and the meso/macroscale physics of volume conduction.

To connect the neural mass and physical frameworks, the first challenge lies in understanding the nature of the current distributions generated by neural activity. A simple approach is to assume that synaptic activity in pyramidal cells generates current dipoles on the cortical surface—a good and useful approximation in the context of EEG measurements given that the electrodes are far from the sources [13]. On the other extreme, local voltage measurements such as LFPs have been modeled using highly realistic models of neurons [14, 15, 16]. The same method has been used to model measurements at different scales [17, 18, 19] with integrate and fire network models. However, this requires a physical model for each cell in the network, making it very demanding computationally. Thus, this approach is difficult to implement at the EEG and SEEG scales and is also not applicable in population based models such as NMMs.

Unlike scalp EEG or LFPs, SEEG measurements fall within an intermediate scale, making it even more challenging to model. On the one hand, SEEG contacts are very close to the current sources, most of the times even surrounded by them, making unsuitable the modeling approaches typically used in the context of EEG (dipole models). On the other hand, the SEEG electrode geometry (contact length: 2 mm, distance between 2 contacts: 1.5 mm) does not permit to have more than one contact implanted within the neocortical thickness. Therefore, the use of highly detailed neuronal models as the current sources in volume conductor models (as is sometimes done in the context of LFP) is technically challenging since they would require the generation of extremely large meshes.

Our first objective is to analyze the current sources in the neocortex at a millimetric scale, adequate for modeling SEEG. For that, we begin at the micro-scale, singlecell level by using highly detailed geometrical models of pyramidal cells together with the multi-compartment cable model formalism [20] to simulate the post synaptic transmembrane currents. We then move to a larger scale and combine single cells into a population to obtain the current distributions that can be expected inside a cortical patch in different situations. These results are then used to simulate SEEG measurements and explore potential simplifications of the current distributions to overcome the challenges associated with the scale of SEEG measurements.

Our second objective is to use the aforementioned modeling framework to create the link between NMMs, which represent the mean activity of neuronal populations, and the cortical CSD associated to this neural activity. The raw outputs of NMMs are the mean synaptic activity, membrane potential alteration and firing rates of each of the populations involved. The mesoscale CSD analysis based on single cell microscale models described above is used to connect the average NMM post-synaptic potentials and the physical model in a straightforward manner and without many assumptions. As a proof of concept of this framework, we used a very simple NMM to generate its associated cortical currents over time and simulate SEEG recordings. We use this example to discuss the limitations of these models in reproducing electrophysiological recordings.

**Figure 1.**
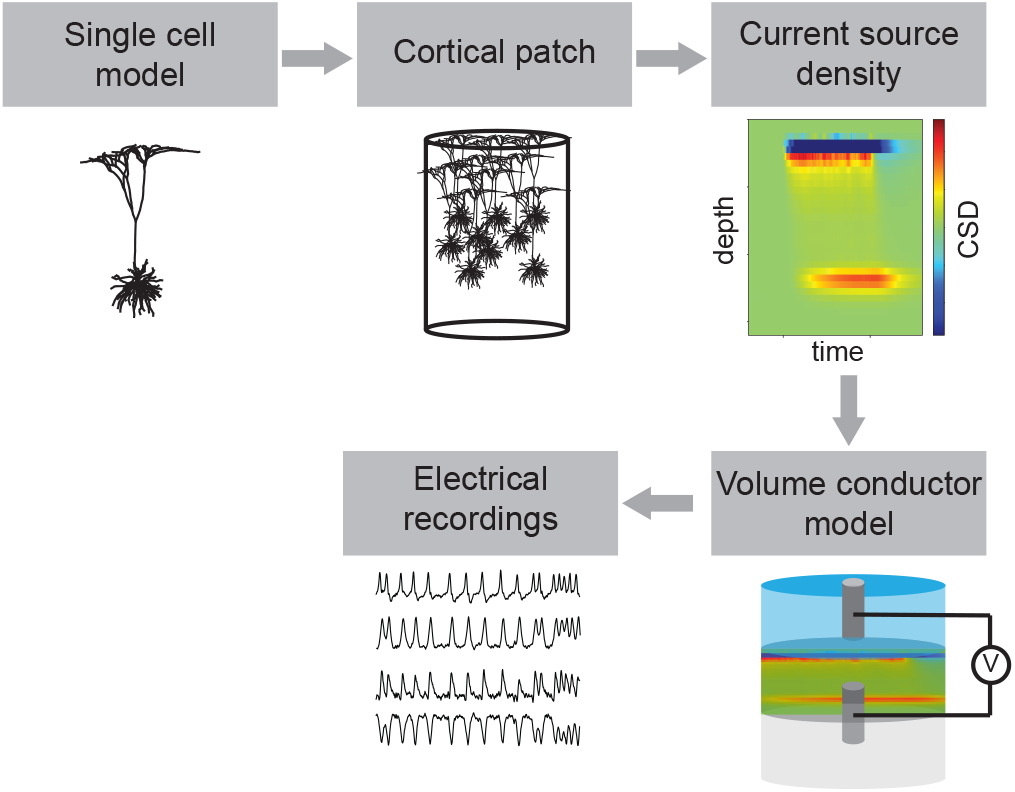
Overview of the multi-scale modeling approach used to generate electrical recordings from highly realistic single cell models.

## 2. Methods

Figure 1 summarizes the modeling approach followed in this study in order to bridge the gap between different scales. First, several (n=1000) highly detailed microscale single cell models are used to generate a cortical patch. The transmembrane currents of these cells are used to calculate the total current inside the cortical patch and obtain an average representation of the CSD depth profile for various scenarios. These CSD profiles can then be used to set the boundary conditions in a volume conductor model for the simulation of electrical recordings. In the present study we did this for a SEEG bipolar measurement (difference between two adjacent contacts), however, the same approach could be used to simulate any recording technique by adapting the volume conductor model. Finally, we propose a link between the neural mass modeling framework and volume conduction physics by taking advantage of the single cell to macroscale recordings scheme implemented.

### 2.1. Single neuron compartmental model

The compartmental cable model formalism [20] was used to simulate the transmembrane currents generated in response of a synaptic input. This was done using LFPy [14, 17], which is an open source Python package for biophysical microscale neural modeling that runs on top of the widely used NEURON simulation environment [21].

The reconstructed cell morphologies of a layer 3 [22], a layer 5 [23] and a layer 6 [24] pyramidal cell from the cat visual cortex were used (see Figure 2a). Since they do not play an important role in the generation of the extracellular signals (we only consider post-synaptic currents in pyramidal cells), the axons were removed from the morphology reconstructions. Layer 3 and layer 6 pyramidal cell geometries were downloaded from `neuromorpho.org` and the geometry of the layer 5 pyramidal cell was downloaded from the LFPY github repository (https://github.com/LFPy). In all cases, the simulations were performed with a passive cell model with the following properties: a membrane capacitance of 1 *μ*F/cm^2^, an axial resistance of 150 Ω cm, and a membrane resistance of 30 kΩcm^2^. The synaptic input was modeled as an exponential decaying current pulse injected through a single compartment with a peak value of *I*_0_=0.65 nA and a time constant of *τ*=2 ms,

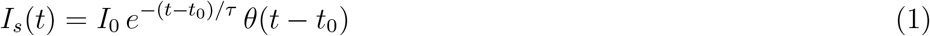

where *θ*(*t*) is the Heaviside step function and *t*_0_ is the peak current time. In all cases, the simulations were run for at least 100 ms before any stimulus with a time step of 1/16 ms.

**Figure 2.**
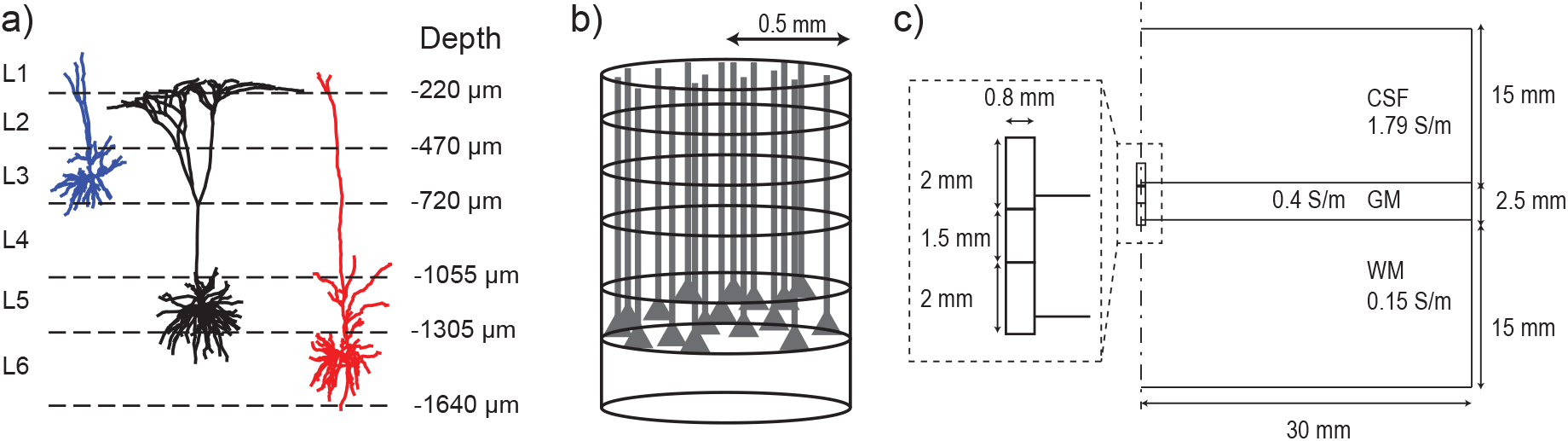
a) Geometries of the pyramidal cells used to generate compartmental models [23, 22, 24]. The figure also shows the different depths at which the cortical layer divisions were set corresponding to the layer size of the cat visual cortex. b) Generation of a cortical patch. Neurons were spatially spread within a 1 mm diameter column with their somas always inside their corresponding layers. The same layer size displayed in a) was used resulting in a GM thickness of 1640 μm. c) Schematic of the axisymmetric geometry used to simulate a pair of SEEG electrode contacts using FEM. Note that the GM thickness in c) is larger than the length of the cortical patch in a) and b). While the latter is meant to represent the average size for humans, the cell morphologies were taken from cat visual cortex. It is also worth noting that the length of an electrode contact is almost the same as the GM thickness.

The synaptic input generates the displacement of charges inside and outside the cell. The injected currents are completed by capacitive return currents in other cell locations.

### 2.2. Cortical patch model

The extracellular voltage that can be measured is the result of the contribution of the voltage generated by the membrane currents of a large number of cells. Thus, in order to create realistic current distributions following a synaptic input we generated a model of a cortical patch (i.e., a cortical column) for each of the three cells described in the previous sections. Our cortical column models model consisted of 1000 neurons (with the same geometries and properties as described in the previous section) placed inside a cylinder with 1 mm diameter. The cylinder was divided along its vertical axis as shown in Figure 2b to replicate the cortical layers. The depths of these divisions can be seen in Figure 2a. These were set to generate a layer size in accordance with the cell morphological models which were all extracted from the cat visual cortex [15].

All cells were perfectly aligned along the vertical axis, but a certain degree of heterogeneity was added by spreading them spatially and rotating them about their vertical axis. For each neuron, a random rotation of [0,2*π*) was applied to the coordinates of its compartments. After this rotation, the center of the soma was placed at a random location inside the cylinder, allowing a vertical spread of ±50 μm around the center of the cell’s corresponding layer. To do so, a random radius [0,0.5] mm and angle [0,2π) were set to define the horizontal position of the soma and a random vertical displacement [-50,50] μm over the center of the layer was set to define the vertical position. Then, a translation was applied to the coordinates of the compartments to place the soma at the desired location.

The overall CSD due to synaptic activity is the result of many neurons receiving many synaptic inputs asynchronously. To replicate that, in our simulations, each neuron received ten pulse inputs at random times each of them injected into a random compartment. The peak current times of these inputs were randomly set with a uniform probability within a time span of 50 ms. Each of these inputs was injected in a compartment that was randomly selected with the function `get_rand_idx_area_norm` from LFPY. This function returns the index of a compartment with a random probability normalized to the membrane area of each segment on the interval [*z_min_, z_max_*]. This same function allowed creating a selection of compartments located exclusively inside a single cortical layer by defining *z_min_* and *z_max_* accordingly. We used this feature in order to simulate the current distribution generated when all cells receive synaptic inputs localized in a single cortical layer.

Once the geometrical and temporal aspects of a simulation were defined as explained above, each cell was simulated individually following the procedure described in section 2.1. Namely, the same passive electrical properties were set, the synaptic inputs were modeled as in (1), and the simulations were run using LFPY.

### 2.3. Calculation of current density and dipole moments

To calculate the current dipole moment over time generated by the transmembrane currents, the center of mass of the current distribution is first calculated at each time point as

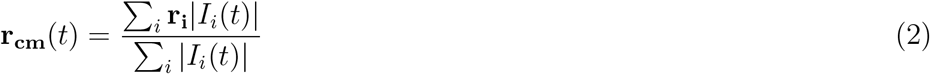

with **r_i_** being the position of the center of compartment *i* and *I_i_*(*t*) the membrane current (ohmnic and capacitive) in compartment *i* at a time *t*. Note that the sum is over all compartments of all cells involved in a simulation. Then, the dipole moment is calculated as

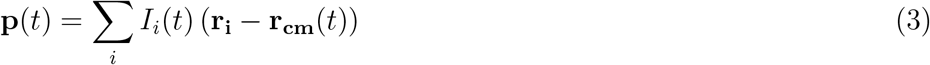

Similarly, the CSD time course was calculated based on the current at each compartment and their positions. To do so, we discretized the cortical patch in 50 μm vertical divisions and summed the membrane currents in the neuronal compartments that were located inside each division. The resulting values were divided by the volume corresponding to these divisions to generate a current density profile over time *CSD*(*z,t*). This volume was calculated as the volume of the disk that is generated after dividing the column vertically (1 mm diameter, 50 *μ*m thick). Note that with this approach we are implicitly averaging the current density over the horizontal directions.

### 2.4. Simulation of SEEG measurements

Next, we used the generated CSD profiles to simulate the voltage that would be measured by what is usually called a bipolar SEEG montage. To achieve this, we built a simple model to simulate the voltage recorded by a pair of two adjacent SEEG contacts belonging to the same lead embedded in a cortical patch. The geometry of the model consisted of three tissue layers to represent white matter (WM), gray matter (GM), and cerebrospinal fluid (CSF). The dimensions of the model and the conductivity assigned to each of these tissues can be found in Figure 2c. We assumed an SEEG lead completely perpendicular to the cortical surface, which allowed us to use an axisymmetric model. SEEG contacts were modeled as 0.8 mm diameter and 2 mm length cylinders separated by an insulating material with 1.5 mm of length [25]. A conductivity of 1000 S/m was set to the contacts and of 10^-5^ S/m to the insulating part between them.

To simulate the voltage difference between the pair of SEEG contacts generated by a certain CSD profile, we took advantage of the reciprocity principle following a similar approach in Moffitt et al. [26]. According to the reciprocity principle, if a unit current is injected through the SEEG contacts and a voltage difference V is measured between two points in space, the same voltage V would be measured between the contacts if a unit current was injected through those two points. In other words, we can calculate the voltage distribution generated when a current is injected by the SEEG contacts and use those results to simulate the voltage measured by those contacts for any given CSD.

The aforementioned approach was implemented using the finite elements method (FEM). The electric potential distribution was calculated by solving the Laplace equation with the geometry described above,

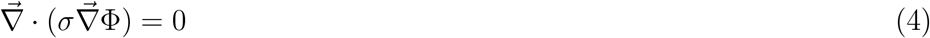

with boundary conditions of a floating potential and a unit ±1 A current (positive on the top electrode and negative on the bottom one) at the contact surfaces. The problem described was solved using COMSOL Multiphysics 5.3a (Stockholm, Sweden) for different depths of the SEEG electrode, *z_el_*. The solutions obtained with this FEM model were used to interpolate the voltage at a grid of points (*ρ, z*) inside the GM. In the radial direction these points were generated every 10 *μ*m beginning at a distance of 100 μm from the surface of the contacts (to account for the fact that scar tissue, and, therefore, no neuronal activity, is expected near the contacts [27]) and extended up to 1 cm away from them. Denser grids of points in the radial direction were tested to ensure that the voltage depth profile did not change. Note that, since an axisymmetric model was used, each of these points is in fact a discrete ring with radius *ρ* around the symmetry axis. Thus, when applying the reciprocity principle the current sources are implicitly modeled as lines (rings) with homogeneous current density.

In the vertical direction, equally spaced points were created to discretize the GM with the same amount of divisions as the CSD profiles in section 2.3. With this approach we re-scaled the CSD profiles to fit the GM dimensions used in the FEM model. Note that the thickness of GM in the FEM model was 2.5 mm to be consistent with realistic human dimensions while the cortical column model from previous sections was 1.6 mm to be consistent with the cat visual cortex, where the neuronal model came from.

The FEM model described above allowed us generating a series of data *V*(*z_el_, r_i_, z_j_*). Then for a given CSD profile, the voltage difference between the SEEG contacts, *V_SEEG_* was calculated using superposition,

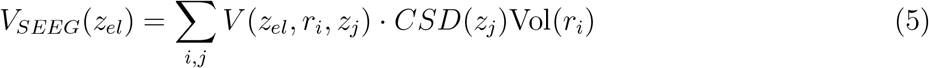

where *V*(*z_el_,r_i_, z_j_*) has units of V/A (voltage when a unit current is injected) and Vol is the volume associated to each point in the grid, which given that we use an axisymmetric model corresponds to the volume associated to a 3D annulus.

### 2.5. Generation of SEEG recordings from Neural mass models

We have used the Jansen-Rit NMM [28] to simulate the average activity of a cortical patch. The Jansen-Rit model includes a population of pyramidal cells (P) that sends and receives inputs from a population of excitatory neurons (Exc) and a population of inhibitory neurons (Inh). The pyramidal population also receives an external excitatory input (Ext). The equations and parameters defining the Jansen-Rit model used for the simulation of cortical activity are described in Appendix A. The parameters of the model were chosen so that the membrane potential of the main pyramidal population displays spikes.

In NMMs, the locations of the synaptic contacts are not explicitly defined — the models are not naturally embedded in space. However, in our framework (as in [10, 11, 12]) they are specified in order to calculate the current distribution that arises from synaptic inputs. For simplicity, we considered the pyramidal cell population in the NMM to be a layer 5 pyramidal cell. Table 1 shows the layer by layer connectivity probability distribution function that was used to locate the synaptic connections to a layer 5 pyramidal cell coming from the other populations in the Jansen-Rit model.

**Table 1.** Probability distribution of the locations of the synaptic contacts from all populations of the Jansen-Rit model into the pyramidal cell population. The table describes the probability for each population to contact each layer.

|  | L1 | L2 | L3 | L4 | L5 | L6 |
| --- | --- | --- | --- | --- | --- | --- |
| External | 2/5 | 2/5 | 0 | 0 | 1/5 | 0 |
| Excitatory | 0 | 0 | 1/6 | 1/6 | 1/3 | 1/3 |
| Inhibitory | 1/8 | 1/8 | 1/4 | 1/4 | 1/8 | 1/8 |

These values provide a coarse representation of a realistic scenario based on the available literature. What follows is a description of the reasoning behind the values selected.

The external input is associated to signals from the thalamus as well as from other cortical regions. In layer 5 pyramidal cells, the thalamic and higher-order cortex inputs mainly contact at the top layers and in a lower proportion at layer 5 [29]. The excitatory population represents the inputs coming from spiny stellate cells and mostly from other pyramidal cells. For simplicity, only the latter were considered. According to Hill et al [30], in layer 5, pyramidal cells seem to make contacts with other pyramidal mostly at the basal dendrites (around 2/3 of the time) and, to a lesser extent, at the apical dendrites (around 1/3 of the time), but not at the tuft. Finally, the inhibitory population represents the inputs coming from several types of Somatostatin expressing cells but, for simplicity, we only considered Martinotti cells. These cells contact approximately half of the time at the apical dendrites and the rest of the time with approximately equal probability at the basal dendrites and the tuft [30]. To use the results from Hill et al, we made the following equivalence: inputs at the tuft are equally distributed between layers 1 and 2, inputs at the apical dendrites are distributed between layers 3 and 4, and basal inputs are distributed between layers 5 and 6.

We used the output of the NMM to calculate what could be regarded as the average post-synaptic current, *I_s_*(*t*), associated to each of the synapses from other populations into pyramidal cells. In fact, we assumed that each of these currents is proportional to the average membrane potential perturbation at each neuron, *u_s_*, induced by the associated pre-synaptic population, *I* = *ηu*, with *η* a gain proportionality factor of units A/V. The gain factor η would correspond to the average current that is required to induce a membrane perturbation of 1 V at the cell soma which depends on many factors (current injection site, cell shape...). For simplicity, we set *η* = 10^-11^ A/mV to obtain equivalent current dipole surface density values in the same order as those measured experimentally which are in the range of 0.16–0.77 nA· m/mm^2^ [31].

In NMMs, *u_s_*(*t*) represents the aggregated or the average effect of a large number of synaptic events at the single cell level. Among other factors, this is proportional to the number of inputs taking place synchronized in time, as well as the average current flowing into the cells at the single cell level for each of these inputs. Thus, *I_s_*(*t*) is a variable that can be linked to the cortical patch model described in the previous section, effectively bridging the gap between microscopic and mesoscopic scales as well as between NMM and the extracellular electrical measurements. To do so, the CSD over time associated to the NMM was calculated as

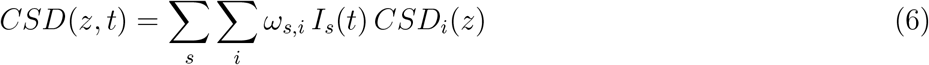

Where *ω_s,i_* is the probability of population *s* to have a synaptic contact in layer *i* (Table 1). *CSD_i_*(*z*) is the CSD profile generated by a unit current input injected at compartments inside layer i to each cell in the patch. The values of *CSD_i_* (*z*) were calculated as described in section 2.3 and were then normalized to a unit current and scaled to the desired cell density as described in the Appendix B. Here we chose a cell density of 100 000 cells/mm^2^ consistent with that usually measured in the cortex [32, 33].

The current source density *CSD*(*z,t*) was then used to calculate the voltage measured by a pair of SEEG contacts as described in section 2.4, as well as the equivalent current dipole moment over time as discussed in Section 2.3 (e.g., for the projection of the NMM activity into a scalp EEG measurement).

## 3. Results

### 3.1. From single neuron to cortical patch

Figure 3a displays the transmembrane current at each compartment at two different time points when an exponential current is injected into a single compartment of the layer 5 pyramidal cell model. Near the peak in the injected current, the return currents are highly localized near the injection site. Later, these return currents spread over the entire cell but the overall current has decreased significantly. Thus, a synaptic input into a single compartment initially generates a very short dipole-like distribution that slowly turns into a longer dipole (but with weaker current) as shown by the CSD profile over time (Figure 3b). These results are in line with earlier modeling [34, 35] and experimental work [36].

**Figure 3.**
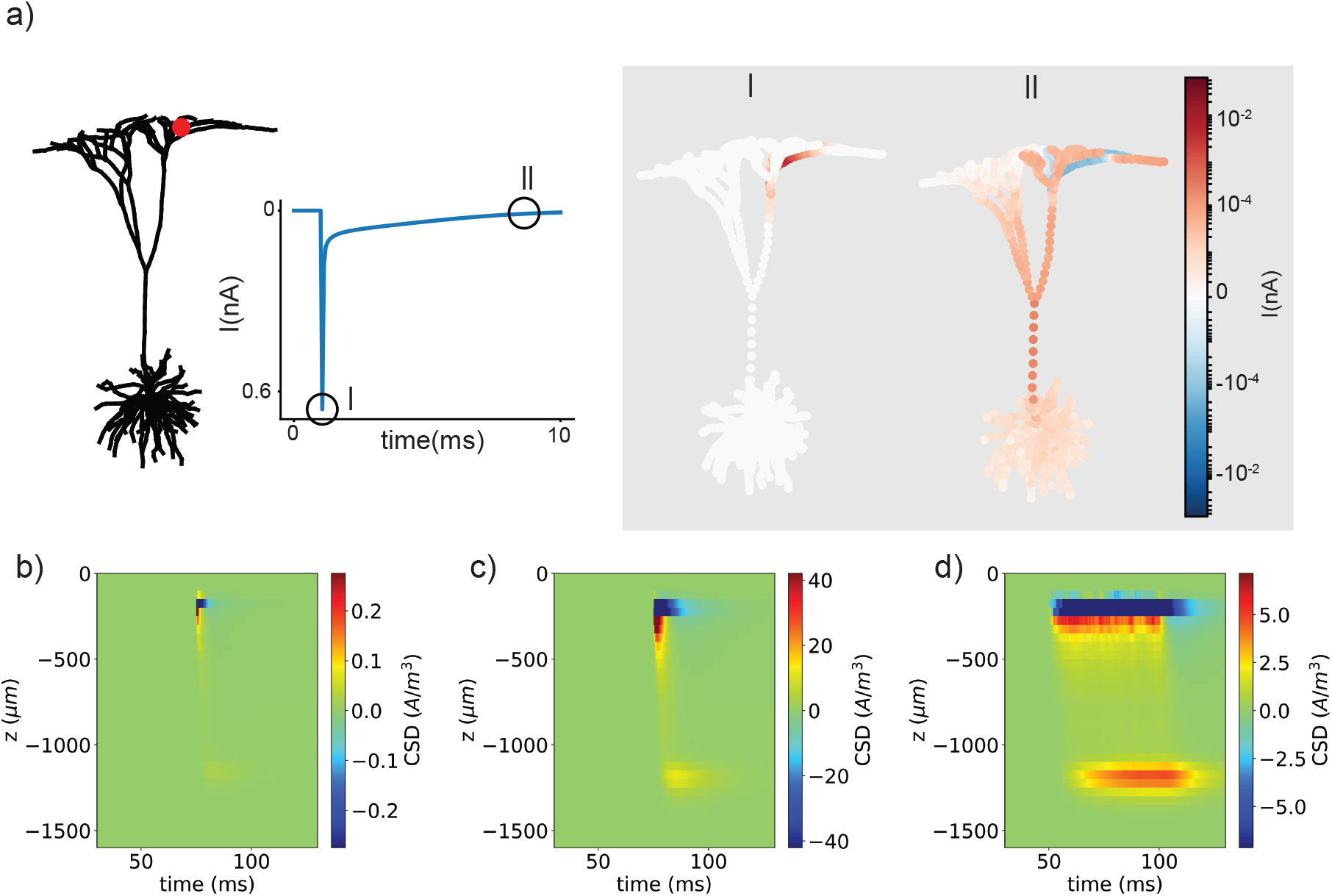
a) Transmembrane current at each compartment in two different time points after the injection of an exponential current into a single compartment of the layer 5 pyramidal cell (marked with a red dot). b) CSD as a function of depth over time for the scenario depicted in a). c) The same plot when one thousands neurons receive synchronously a single input in a compartment located in layer 1. d) The same plot when these neurons receive the input asynchronously into compartments located in layer 1. It is important to note that all color scales in b-d were adjusted so as their limits were 1/3 of the maximum value for a fair comparison.

Figure 3c provides depth-time plots of the CSD obtained when an input is injected synchronously at the same location into one thousand cells located inside a cortical patch (see section 2.2), with the input injected into a random compartment located in layer 1. The CSD over time shows a similar pattern than the single cell case, but the magnitude of the CSD during the initial short dipole and the later long dipole are more similar. In fact, a few milliseconds after the injection, the return currents around the location of the soma generate a CSD that is comparable to the CSD generated near the injection site at the peak in current. This is due to the spatial summation of the currents from various cells spread in space. Right after the peak in injected current, the current is highly localized near the injection site, and the currents from different cells largely cancel each other out. However, later, when the return currents spread over the entire cell, the currents from individual cells add in a much more efficient way.

The effect described above is further reinforced in a more realistic scenario, with each cell receiving the input at a random time within a 50 ms time window (Figure 3d). Because of the spatiotemporal smoothing induced by the cable equation, both the spatial input spread and the input time randomization contribute to a longer lasting dipole-like temporal distribution.

### 3.2. Synaptic inputs at different layers

We simulated a cortical patch (see section 2.2) in which each neuron receives ten synaptic inputs at a random time within a 50 ms time window. These inputs were injected at a random compartment of the cell belonging to a specific layer in order to analyze the CSD depth profiles that are generated depending on the location of the synapses. These results, displayed in Figure 4, were generated for each cell model individually. Figure 4a provides the results obtained for layer 3 pyramidal cells receiving inputs at different layers. The same results for layer 5 and layer 6 pyramidal cells are shown in Figure 4b and Figure 4c respectively. Note that for layer 3 pyramidal cells inputs were simulated only in layers 1-3 because there are no compartments present in deeper layers.

**Figure 4.**
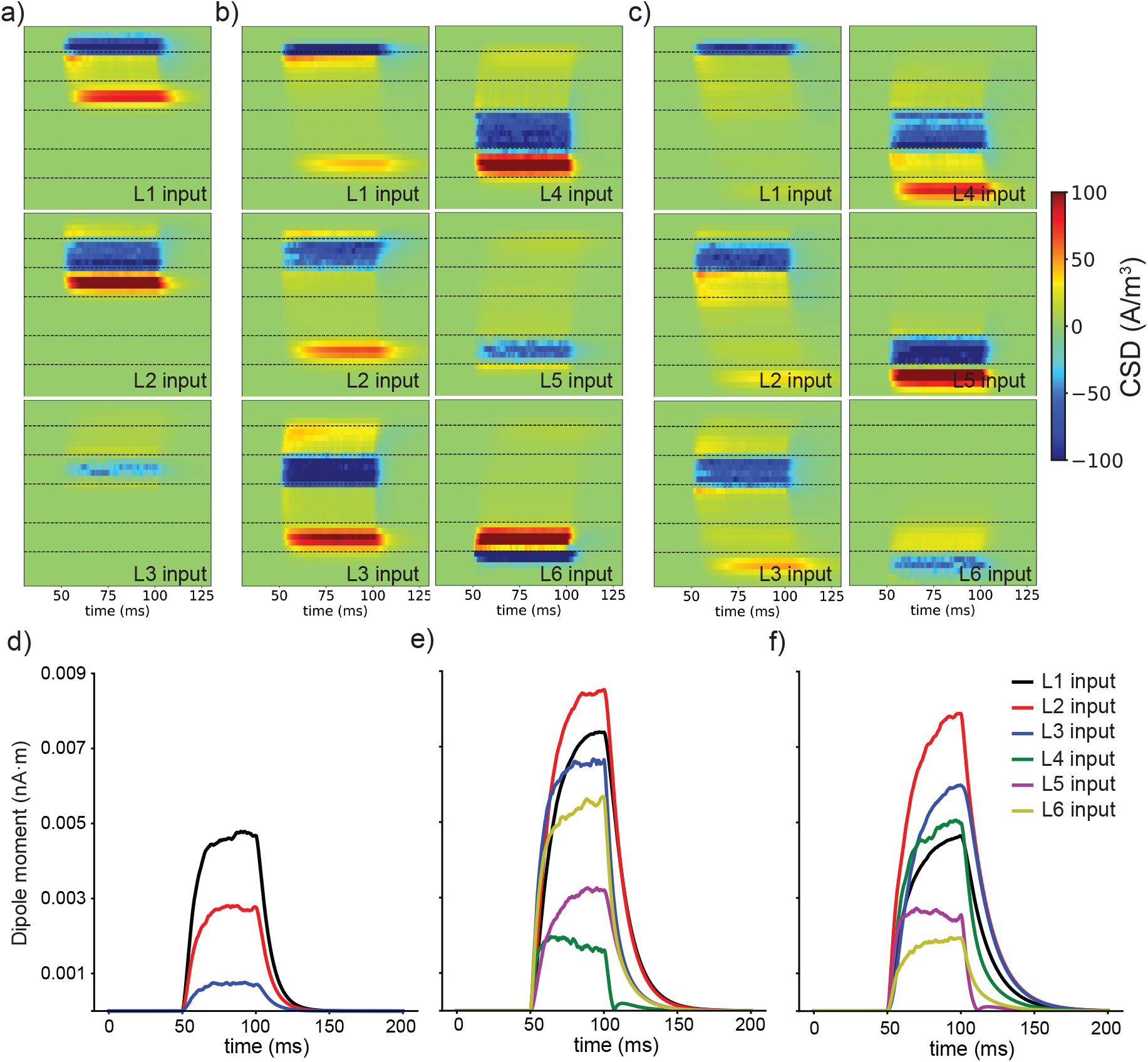
a) CSD as a function of depth (vertical axis) due to synaptic inputs limited to only one of the layers in a cortical patch containing layer 3 pyramidal cells. The three rows provide results for different input layers (see embedded legend). b) The same results for a patch of layer 5 pyramidal cells and c) layer 6 pyramidal cells. The dashed lines represent the limits of the cortical layers. Current dipole moments over time generated by the previous current distributions for layer 3 (d), layer 5 (e) and layer 6 (f) pyramidal cells.

As expected, the negative current in these CSD distributions is exclusively located in the compartment where the inputs are injected. Interestingly, a very similar pattern is observed for all three cell models. In all cases, a large portion of the return current is localized at the layer where the cell soma is located, except when the input targets the soma directly. This is is due to the contribution of all the basal dendrites that are mostly located there and constitute a large portion of the cell surface. A similar but milder effect can be seen in some of the cases at the top layers due to the presence of a large number of branches of the apical dendrites. When the input is localized in the layer where the soma is placed, a large portion of the return currents is confined into the same layer. This results in a large cancellation of the currents as evidenced by the fact that those cases display the smallest current values.

It is important to note that, although none of the obtained CSD profiles have a perfect dipole-like distribution, they all resemble a dipole to a certain extent. However, the strength of these dipoles changes depending on the layer at which the synapses are located. Depending on their location (Figure 4d-f), synaptic inputs can generate dipoles differing by a large magnitude (up to four times). Specifically, inputs at the top layers, above the soma, generate dipoles that are at least two times stronger that those generated by inputs at (or close to) the soma layer. This is due to 1) the larger distance between sources and sinks and 2) the large cancellation that takes place for inputs at the soma layer discussed above.The difference between the top layers (1-2) and the soma layers is particularly relevant since they contain a large portion of the dendritic arbor making synaptic contacts there more likely.

### 3.3. Simulation of SEEG measurements

The CSD profiles obtained for the inputs at different layers were used to simulate an SEEG measurement (see Section 2.4). The results in Figures 4a–c were averaged over the time window in which the dipole moment is larger (70–100 ms) and rescaled to fit human cortical dimensions. The resulting CSD profiles as a function of depth are presented in Figure 5a. These, together with a FEM model, allowed us to obtain the voltage difference that a pair of SEEG contacts would measure. Figure 5b displays the voltage difference measured between two consecutive contacts of an SEEG lead for different depths and for the different locations of synaptic inputs and the three cell models.

**Figure 5.**
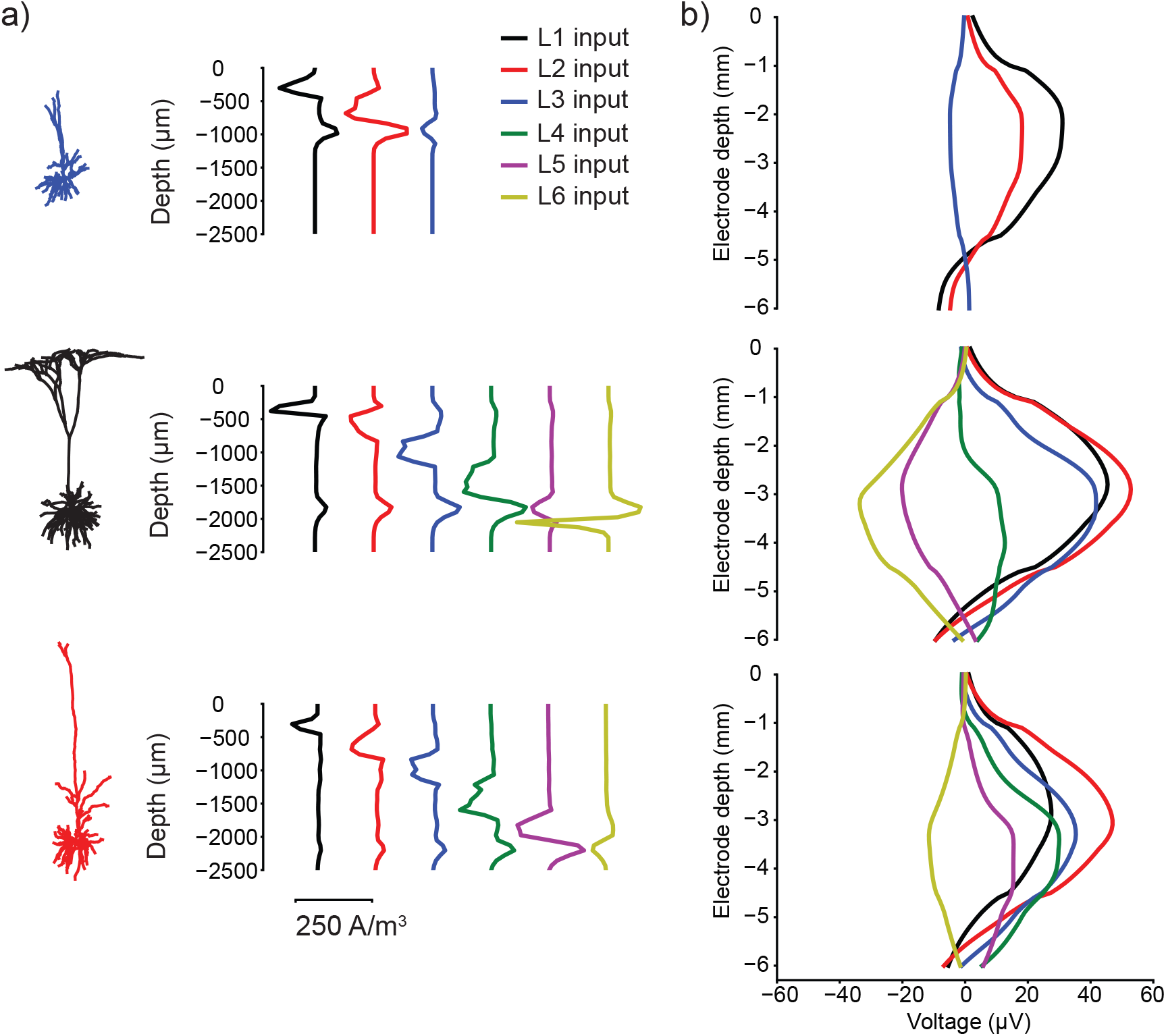
a) CSD profiles generated by synaptic inputs at different layers in the three cell models. The results were re-scaled to fit the human cortical size. b) Bipolar voltage difference between two consecutive SEEG contacts at different depths when they are surrounded by the different CSD profiles in a). Electrode depth origin is set at the location in which the deepest contact has its center aligned with the GM/CSF boundary. At a depth of −6 mm, the outermost contact has its center aligned with the GM/WM boundary. The results are shown for layer 3, layer 5 and layer 6 pyramidal cells (from top to bottom).

From the diversity of CSD profiles in Figure 5a it is clear that the voltage difference depends on the location of the synaptic inputs (as will the equivalent dipole current strengths of these distributions). Associated with this, there is a large variation in the potential difference depending on the electrode depth since the relative positions of the sinks/sources and the electrodes drastically change as the electrode is moved (Figure 5b). However, a remarkable observation is that the relative magnitude of the voltage difference for inputs at different layers changes with electrode depth. This implies that the relative contribution of each synaptic input over the measured voltage depends on the electrode depth, making SEEG recordings highly sensitive to electrode depth.

### 3.4. Simplified CSD model for SEEG

We then compared these voltage measurements with those obtained with the same FEM model but using a much simpler current source model consisting of arrays of discrete sources placed at the center of each layer. For each cell and synapse position (synapse connecting to each of the layers) we generated a simplified model that was heuristically adjusted to fit the results presented in Figure 5b. In order to simplify as much as possible the CSD representation, for each case, only a source and a sink location were allowed. Figure 6 shows the distribution of the currents on these simplified models. Note that the magnitude of the discrete sources was normalized to their maximum value. The same figure provides the voltage difference measured by a pair of SEEG contacts (obtained from the same FEM model) in comparison with that obtained with the sources modeled from the compartmental models (Figure 5a). The voltage profiles generated with the simplified models replicate quite well those obtained with the original CSD.

**Figure 6.**
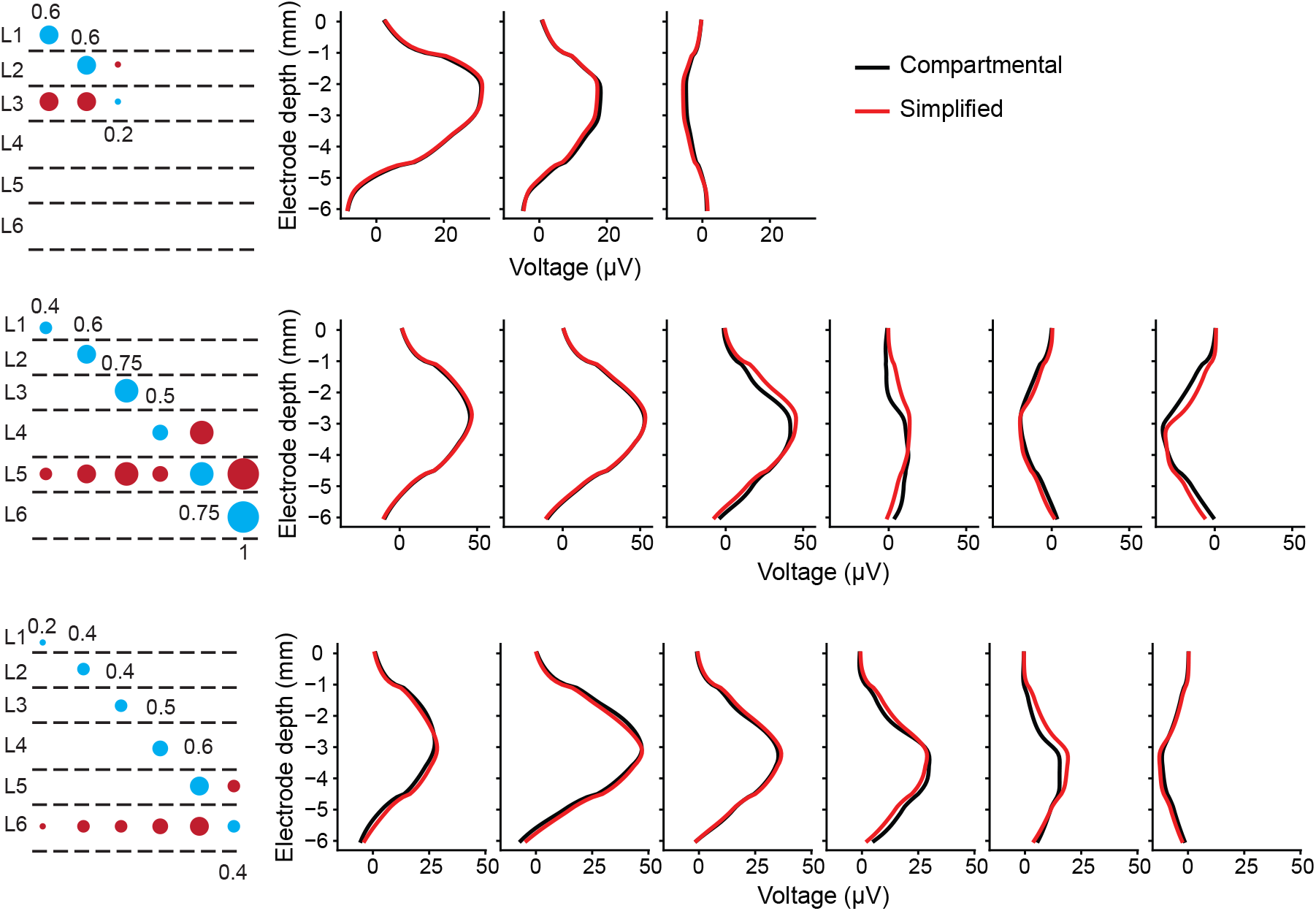
On the left, simplified current sources created to replicate the voltage profiles in Figure 5b. The graphs display the locations and values of the discrete current sinks (blue circles) and sources (red circles). The size of the circles and the numbers next to them correspond to the current strength relative to the strongest one. On the right, comparison of the voltage profiles obtained with the CSD profiles in Figure 5a and those obtained with the simplified model. The results are shown for layer 3, layer 5 and layer 6 pyramidal cells (from top to bottom).

### 3.5. Voltage recordings from NMM

Figure 7a provides a schematic of the distribution of the synaptic inputs into the pyramidal cell of the Jansen-Rit model. These distributions, together with the results in Figure 4b, were used to calculate the CSD over time produced by the neural activity in the Jansen-Rit model (Figure 7b). Sinks and sources can originate from a synaptic input or by the return currents associated with the input. This makes their interpretation ambiguous even in a simple scenario such as the one depicted here (only three synapses and one type of pyramidal populations). In layers 1–2, sinks can be either due to the external noise input or the return currents associated to the inhibitory inputs while the sources in this model can only be associated to the return currents in the excitatory population. In layers 3–4, practically no return current is expected. Therefore, sources in those layers can only be due to the inhibitory inputs. Sinks associated with the excitatory inputs are also generated in these layers, but they do not appear in the CSD plot because they are masked by the inhibitory inputs. Layer 5 constitutes the most ambiguous region since all populations target in some proportion the dendrites there, and, at the same time, a large portion of the return currents are expected to take place in this layer. In particular, sources can be due to inhibitory inputs or the return currents of the other input types, while sinks can be due to external and excitatory inputs or the return currents from inhibitory inputs. Finally, in layer 6 the sinks related to the excitatory input dominate the CSD. Sources due to the inhibitory inputs are also present in this layer but they are hidden by the sinks.

**Figure 7.**
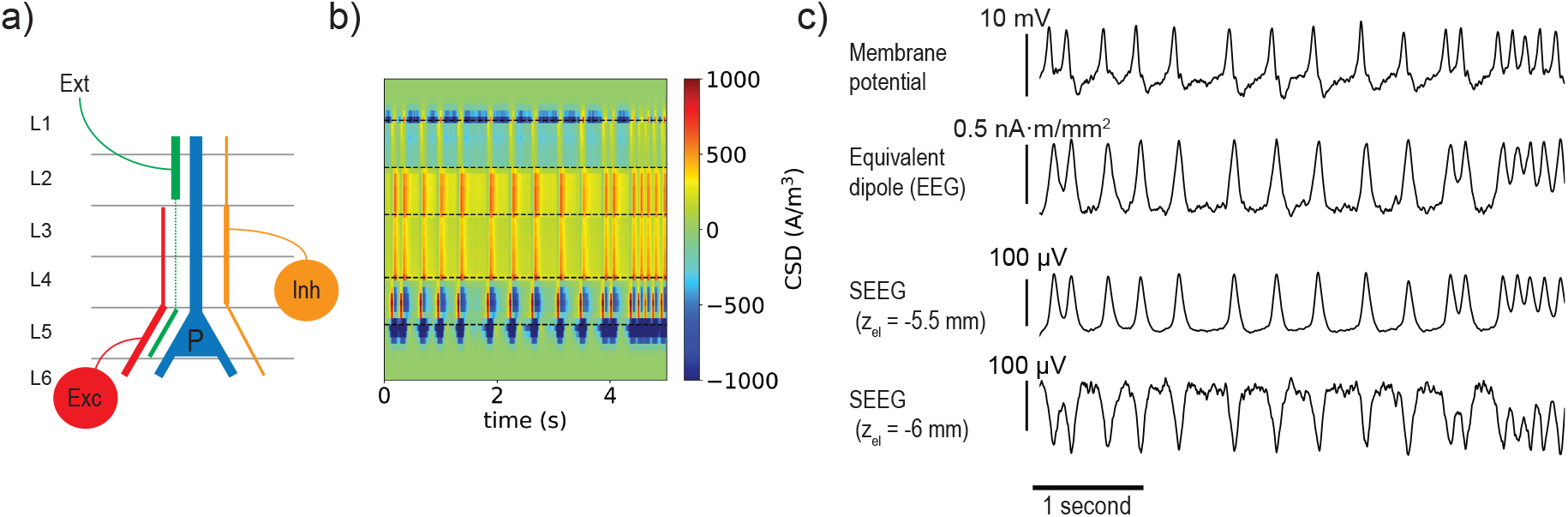
a) Schematic of the probability distribution of the synaptic contacts on the pyramidal cell for the different inputs. The thickness of the lines is proportional to the probability of contact (see Section 2.5). Note that the positions of the circles that represent the inhibitory and excitatory populations are not meant to represent the locations of the cells belonging to them. b) CSD as a function of depth over time obtained with the Jansen-Rit model and the synapse distribution in a). c) Electrical recordings generated with the Jansen-Rit model using different signal proxies: the pyramidal cell membrane potential from the NMM, the equivalent current dipole of the CSD (i.e. what would be projected to a scalp EEG measurement) and the properly simulated bipolar SEEG at different depths (last two graphs).

The simulated CSD can be used to generate SEEG signals and compare them with different proxies sometimes used in the field. Figure 7c displays the membrane potential of the simulated pyramidal cell population. This is the sum of all the post-synaptic membrane perturbations and is commonly used directly as a proxy for both scalp and intracranial EEG measurements. The figure also shows the equivalent current dipole moment of the CSD over time, which is essentially the waveform that would be recorded by a scalp EEG electrode from this cortical patch. Finally, the SEEG simulated recordings for two different depths are also displayed.

Even though the same patterns or event types can be recognized in the four traces, they all look slightly different. Namely, in all four traces we can observe spikes happening with the same frequency, but the shape of these spikes can change. While in the NMM membrane potential most spikes display a positive peak followed by a negative smaller peak, in the other traces a first peak is usually followed by a return to baseline. Another noticeable difference between waveforms is that the contribution from the external input (higher frequency ripple) is more visible in some of them than the others. It is also important to note that with a small displacement of the electrode (hundreds of μm), the SEEG recorded waveform can substantially change. Here we selected as example two positions with 500 μm distance between them displaying a change in polarity and a difference in the shape of the waveforms recorded. From Figure 5, it can be deduced that the same degree of variation can be expected for many other positions.

## 4. Discussion

### 4.1. Validity of the results

The flow of ionic charges that generate the electrical fields measured by SEEG or EEG is the result of the temporal and spatial summation of a large number of synaptic inputs over a large number of cells. This is nicely illustrated in Figure 3, which highlights the importance of considering both the temporal and the spatial aspect of this summation when generating the current distribution.

In order to compute the different CSD profiles, in this paper we modeled a scenario with one thousand cells each receiving ten inputs within the same time window (corresponding to an average input frequency of 200 Hz). While these numbers are rather arbitrary, they are large enough to ensure that the results would not change qualitatively by increasing them. That is, given that the superposition principle would apply, our simulations are in a limit in which increasing the number of inputs or the number of cells changes the magnitude of the CSD but the overall depth profile pattern remains the same. The density of pyramidal cells in the cortex is usually in the order of 100 000 cells/mm^2^ [32, 33], which is two orders of magnitude higher than in our simulations. Thus, it is very likely that in a realistic setting every cortical patch operates in this regime, i.e., where the CSD magnitude depends on the number of synaptic inputs synchronized in time but its overall profile does not.

Measurements in humans indicate that current dipole surface densities in the neocortex are in the range of 0.16–0.77 nA· m/mm^2^ [31]. This is two orders of magnitude higher than our results (see Figure 4d-f), which is in fact consistent with the two orders of magnitude difference in cell density between our simulations and what is measured experimentally. In contrast, despite of the cell density, we obtained voltage differences only one order of magnitude lower than those measured experimentally, which is in the hundreds of *μ*V (see [37]). This could be due to 1) the synaptic input current we used being too high (which is unlikely given that dipole strength is consistent with experimental data), or 2) the axisymmetric geometry of our model excessively favoring the spatial summation of the current sources—our model assumes that the SEEG contacts are surrounded by a 1 cm patch of perfectly aligned sources with a homogeneous activity, or 3) the cell density in our model being too concentrated in specific layers, which will impact local electric field but not overall dipole.

It may be argued that since throughout the entire study only one cell morphology was used to represent each pyramidal population, our results may be biased due to the model choices. However, while there may be a certain bias in quantitative terms, we do not expect our results to be different qualitatively if more geometrical representations of the same cell populations were used. In fact, the CSD profiles in Figures 4a–c display very similar patterns (return currents mostly confined to the regions near the soma), reinforcing the idea that our results in qualitative terms are not biased due to the choice of the cell morphologies.

### 4.2. Brain current sources and SEEG volume conduction models

The CSD profiles that were obtained for the different synaptic input locations (Figure 4b) can all be regarded as dipole-like current distributions to some extent. However, the center and strength of their equivalent dipoles are very different among them. Thus, unlike it is commonly done in EEG, all these synaptic input types cannot be modeled with a single dipole to reproduce SEEG voltage measurements. However, using a set of dipole-like distributions we were able to reproduce the voltage profiles for single input types measured by a pair of SEEG contacts quite accurately (Figure 6). Therefore, the simplest model that could be used to represent the brain sources in the context of SEEG would need to include six current-conserving monopolar sources (one per layer). This could be a useful approach if SEEG measurements were to be modeled in more complex volume conductor geometry than the axisymmetric model used in this study. In that case, the use of such simplified model would help reducing the mesh resolution required to set the boundary conditions required to reproduce the CSD.

Our results also show that the location of the synaptic contacts play a critical role in shaping the current distributions generated by different types of brain activity. This has implications in the context of SEEG (Figure 5b) but also in the context of EEG since the dipole strength can vary significantly (Figures 4d–f). As already discussed in other studies [18], this implies that the different pre-synaptic populations do not contribute equally to the equivalent cortical dipole that they generate. Thus, although scalp measurements are taken far enough that the CSD profile can be regarded as a current dipole, the strength of this dipole is the result of a weighted sum of the different types of post-synaptic currents. These weights will depend on the locations of the synaptic contacts. In this study we proposed a modeling framework that takes this fact into account and can be implemented to simulate both SEEG and scalp EEG recordings. We created an example of this approach as a proof of concept (Figure 7). In the next section we discuss the validity of this approach and the implications of our results.

### 4.3. Link between NMM formalism and physical measurements

The discussion in Section 4.1 is also relevant to the link between the NMM formalism and the volume conduction physics proposed in this study. If we assume that a cortical patch always operates in a regime in which the CSD profile generated by a certain type of neural activity is independent of the amount of synaptic events (CSD magnitude is scaled by the amount of inputs, but its shape does not change), then the link between NMM activity and the corresponding CSD over time is straightforward. NMMs are meant to represent the average activity of interconnected neural populations over a large number of cells. This is a mathematical abstraction, but the variables that describe the dynamics in these models can be used as a starting point for a physical model. The firing rates and membrane potential perturbations can be interpreted as a measure of the degree of synchronization of the neurons within a population inside a cortical patch. Thus, the post-synaptic potentials in the pyramidal cells can be regarded a measure of the average amount and strenght of the synaptic inputs (i.e., average post synaptic currents) that these cells receive over time. With this interpretation, the magnitude of the CSD profiles are proportional to the post-synaptic currents associated to each population but the location of the sources and sinks is always the same. This approach nicely links the microscopic (single neuron and single input) and mesoscopic (large number of cells and inputs) scales without any unreasonable assumptions.

In the present study, we based all the analysis of the CSD profiles on the classification of synaptic inputs with respect to the cortical layers they target. With this approach, we obtained CSD profiles that resemble those measured experimentally (sinks and sources bounds seem to match the cortical layer divisions) [38, 39]. But more importantly, by doing so, we provided a framework to connect NMMs with a physical model that can be used to simulate any type of electrical recordings. Although we discussed it in the context of a Jansen-Rit model, our approach can be generalized to a wide range of NMM simply by adjusting the locations of the synapses within the pyramidal cell populations. That is, each population in a NMM represents a certain set of cell types, and for each cell type it is possible to define a distribution of the synaptic contacts into the pyramidal cells based on experimental studies (some examples can be found elsewhere [40, 41, 29, 30]). This allows modeling the contribution to the CSD profile of each population in the NMM, which can be used to calculate the CSD profile over time generated by the NMM activity. Once the CSD is calculated, it can be used to set the boundary conditions of a volume conductor model and simulate any type of electrical measurements following the same approach as we did here for an SEEG measurement.

The membrane potential of the pyramidal cell population taken directly from NMMs is sometimes used in the literature as a proxy for scalp EEG or SEEG recordings. This implies assuming that the membrane potential is equivalent to the cortical current dipole strength (in scalp EEG) or completely ignoring the volume conduction side of the problem (in the case of SEEG). We have shown that even in a simple NMM, the membrane potential and the equivalent current dipole display different waveforms. This is especially relevant when simulating SEEG recordings since, as we showed, the recorded waveform is highly sensitive to the electrode contact locations with respect to the current sources. However, it is important to note that, in essence, the four waveforms in Figure 7c are a weighted sum of the same signals. The membrane potential from the NMM implicitly assigns the same weight to all post-synaptic currents, while in the other cases the contribution of each of them is different (it depends on the locations of the synaptic contacts of each population, the volume conduction model and the electrode locations). In fact, earlier work has already pointed out that that a weighted sum of post-synaptic currents is a good proxy for EEG signals generated with a leaky integrate and fire neural network model coupled with highly realistic volume conduction models [18].

In summary, to address the aforementioned issue and simulate electrical recordings with NMMs more realistically from a physics viewpoint, one should:

- Define the distribution of the inputs for each pre-synaptic population on the pyramidal cells.
- Calculate the CSD profile over time from the NMM based on the synaptic input distribution.
- Generate an adequate volume conductor model by taking into account the scale of the measurements that are simulated.
- Use the volume conductor model to project CSD over time into electrode voltage measurements (usually known as the forward problem).

Because of the principle of superposition (linearity), from this modeling framework some elementary conclusions can be drawn about the general spectral content of the electrophysiological recordings simulated with NMMs. First of all, if a frequency (or oscillatory pattern) is not present in any of the post-synaptic currents (i.e., none of the activities of the neural populations oscillates at that frequency), it will not appear in electrophysiological measurements regardless of the recording technique or electrode positions. Second, although in SEEG the recorded signal depends on the position of the contacts, the frequency content is always the same, only the power spectral density shape can change. Related to that, if two frequencies are only present in one of the post-synaptic currents, then the energy ratio between them is maintained regardless of the recording position.

All these ideas may help to set the limits of what features of the electrical recordings can be aimed to be reproduced using NMMs without a proper physical model. While the oscillatory patterns or chains of events (e.g. the stages of an epileptic seizure) will appear in the simulated waveforms regardless of the volume conductor model, the shape of the waveforms can only be simulated by a volume conductor model. Similarly, great care is required when interpreting experimental waveform shapes (e.g., energy ratio between frequency bands or complex spike morphologies), since, in addition to physiology, they will strongly depend on the location of synapses and the physical environment of the measurements.

## 5. Conclusions

Starting from the model of synaptic and return currents associated with the realistic representation of a single neuron and some basic assumptions on the distribution of neurons (vertical distribution of synapses, horizontal isotropy), we simulated the activity of a cortical patch to bridge the gap between the microscale and mesoscale levels. The model generates CSD profiles that can be combined to represent different brain activity patterns at the mesoscale level. These profiles can be used together with a volume conductor model to simulate SEEG, scalp EEG or other types of electrical recordings. However, since they were generated with a computationally-demanding resolution, we also investigated simplified versions that produce essentially the same results after parameter fitting. We showed that with adapted discrete monopolar sources it is possible to reproduce the voltage measured by a pair of SEEG contacts essentially in the same way as with the more detailed model.

Our cortical patch model provides a connection between the NMM formalism and the physics of electrical measurements. We showed that when the geometry (locations of the synaptic contacts) and the physics (volume conduction model, electrode geometry) are taken into account, the post-synaptic currents in pyramidal cells contribute differently to the recorded electrophysiological signals. This is especially relevant when modeling SEEG, as the contributions vary significantly depending on the position of the electrode. Finally, we also showed that, for these reasons, proxies such as the membrane potential of the pyramidal population in a NMM can be misleading, resulting in distorted EEG spectral and waveform shapes. Our modeling framework overcomes these limitations and can be generalized to arbitrary laminar NMMs implemented into appropriate volume conductor models and measuring electrode configurations.

## Appendix A. Description of the Jansen-Rit neural mass model

The NMM is characterized by state variables describing the firing rate of each neuronal population and the average membrane perturbations induced by each synapse.

The average membrane potential *v_n_* of the cells in the population is the summation of all the average pre-synaptic membrane perturbations *u_s_*, and its average firing rate *φ_n_* is computed using a non-linear function of the membrane potential,

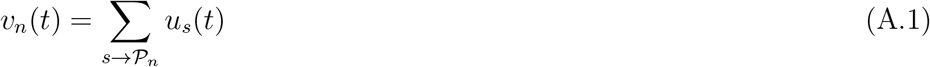

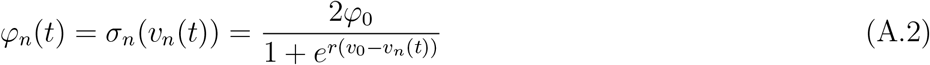

where the sum in the first equation is over synapses reaching the population, *φ*_0_ is half of the maximum firing rate of each neuronal population, *ν*_0_ is the value of the potential when the firing rate is *φ*_0_ and *r* determines the slope of the sigmoid at the central symmetry point (*ν*_0_, *φ*_0_) [28].

Each synapse *s* is described by an equation representing the conversion from an input firing rate *φ_n_* from the pre-synaptic population *n* into an alteration of the membrane potential *u_s_* of the post-synaptic neuron. This relation is represented by the integral operator 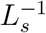 (a linear temporal filter), the inverse of which is a differential operator *L_s_*,

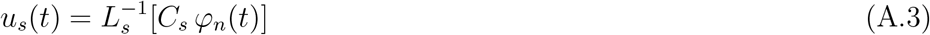

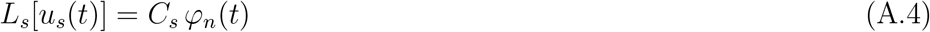

where *C_s_* is the connectivity constant between the populations. The differential operator *L_s_* is defined as

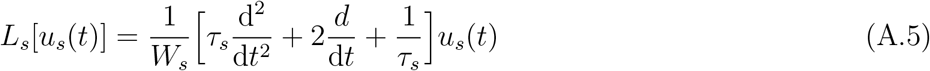

where *W_s_* is the average excitatory/inhibitory synaptic gain and *τ_s_* the synaptic time constant. This time constant is a lumped parameter representing the population averaged effective delay and filtering time associated with the time from reception of input in the cell to soma potential perturbation.

We simulated 5 seconds of neural activity using the model described here and the parameters listed in Table A1. The model equations were integrated using a 4th order Runge-Kutta scheme and a time step of 1 ms.

## Appendix B. CSD re-scaling

In the NMM formalism, the output magnitudes are in terms of average per cell in each population. In section 2.5 we explained how do we obtain the average post-synaptic current per cell in the pyramidal cell population, *I_s_*. In order to connect these values with the CSD profiles generated we need to first normalize them to an average unit input current (1 A) and a unit cell density (1 cell/mm^2^).

**Table A1.**
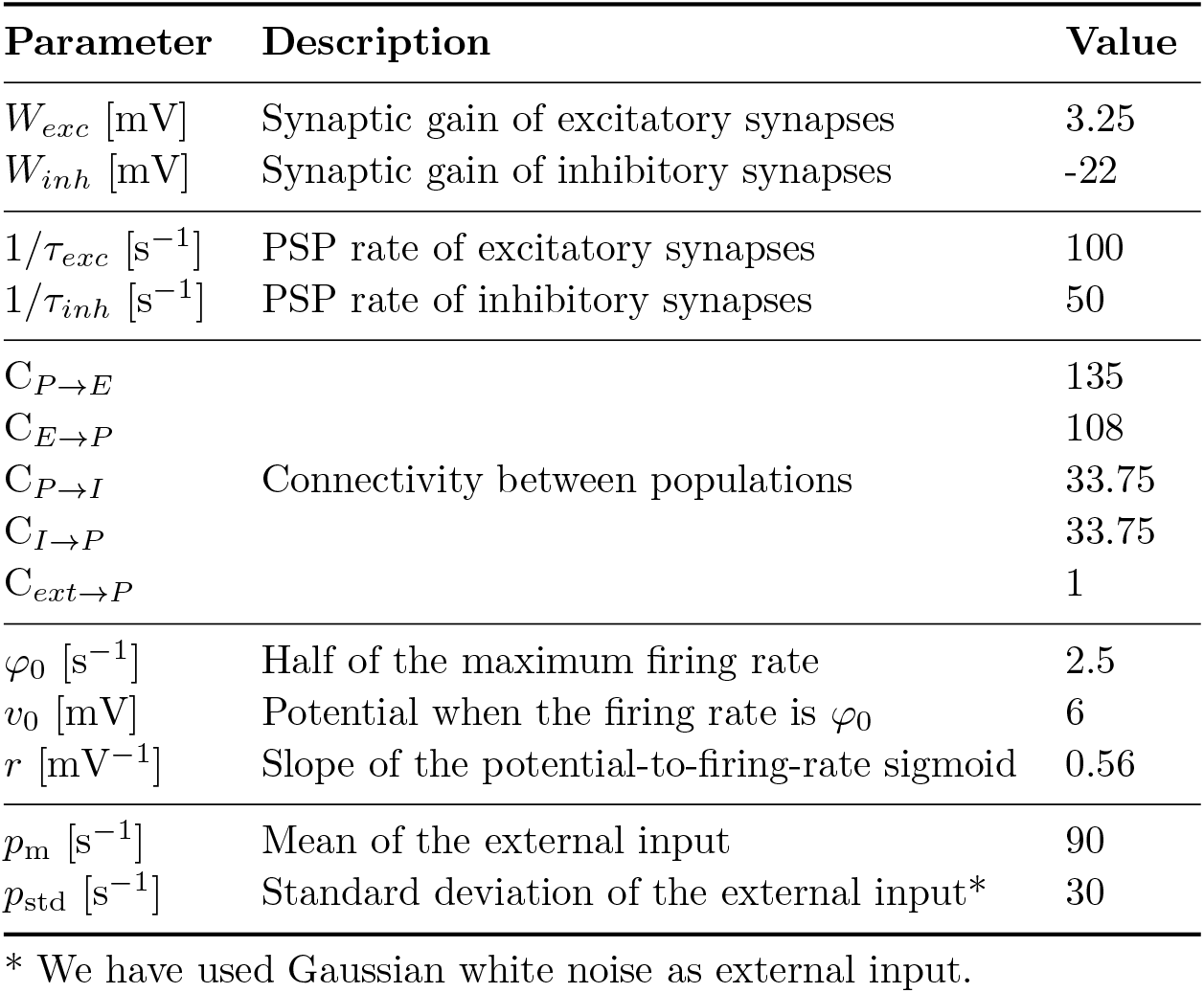
Parameters of the Jansen-Rit NMM used for the simulations.

The current inputs injected are exponential decays of the form *I*_0_*e*^-*t/τ*^. Thus, the charge injected per input is:

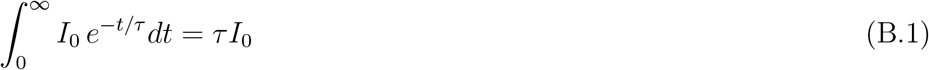

Considering that 10 inputs are injected per cell within a 50 ms time window, the average current injected per cell is:

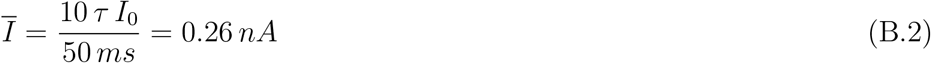

We can then use 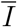 to re-scale our CSD profiles to a unit synaptic input current. Similarly, we can re-scale the CSD profiles for any cell density. Our simulations contained 1000 cells in a 0.5 mm diameter cylinder which amounts to a density of around 1273 cells/mm^2^. With these values, we can calculate the CSD that would correspond to a single cell inside a 1 mm^2^ patch receiving a 1 A average current input. Once these normalized CSD values have been obtained, we can re-scale them to the desired cell density and average input currents.

## Acknowledgments

This work has received funding from the European Research Council (ERC) under the European Union’s Horizon 2020 research and innovation programme (grant agreement No 855109; ERC-SyG 2019) and from FET under the European Union’s Horizon 2020 research and innovation programme (grant agreement No 101017716)

## References

[1] Fabrice Bartolomei, Anca Nica, Maria Paola Valenti-Hirsch, Claude Adam, and Marie Denuelle. Interpretation of SEEG recordings. Neurophysiologie Clinique, 48(1):53–57, 2018.

[2] György Buzsáki, Costas A. Anastassiou, and Christof Koch. The origin of extracellular fields and currents-EEG, ECoG, LFP and spikes. Nature Reviews Neuroscience, 13(6):407–420, 2012.

[3] V. K. Jirsa, T. Proix, D. Perdikis, M. M. Woodman, H. Wang, C. Bernard, C. Bénar, P. Chauvel, F. Bartolomei, F. Bartolomei, M. Guye, J. Gonzalez-Martinez, and P. Chauvel. The Virtual Epileptic Patient: Individualized whole-brain models of epilepsy spread. NeuroImage, 145:377–388, 2017.

[4] Fabrice Wendling, F. Bartolomei, J. J. Bellanger, and P. Chauvel. Epileptic fast activity can be explained by a model of impaired GABAergic dendritic inhibition. European Journal of Neuroscience, 15(9):1499–1508, 2002.

[5] F.H. Lopes da Silva, A. van Rotterdam, P. Barts, E. van Heusden, and W. Burr. Models of neuronal populations: The basic mechanisms of rhythmicity. Progress in Brain Research, pages 281–308, 1976.

[6] FH Lopes da Silva, A Hoek, H Smits, and LH (1974) Kyber-netik 15: Zetterberg. Model of brain rhythmic activity: the alpha rhythm of the thalamus. Kybernetik, 15(1):27–37, 1974.

[7] A van Rotterdam, F H Lopes da Silva, J van den Ende, M A Viergever, and A J Hermans. A model of the spatial-temporal characteristics of the alpha rhythm. Bulletin of Mathematical Biology, 44(2):283–305, 1982.

[8] Behnam Molaee-Ardekani, Javier Márquez-Ruiz, Isabelle Merlet, Rocio Leal-Campanario, Agnès Gruart, Raudel Sánchez-Campusano, Gwenael Birot, Giulio Ruffini, Josá-Maria Delgado-García, and Fabrice Wendling. Effects of transcranial direct current stimulation (tDCS) on cortical activity: a computational modeling study. Brain Stimul., 6(1):25–39, January 2013.

[9] Isabelle Merlet, Gwénaël Birot, Ricardo Salvador, Behnam Molaee-Ardekani, Abeye Mekonnen, Aureli Soria-Frish, Giulio Ruffini, Pedro C Miranda, and Fabrice Wendling. From oscillatory transcranial current stimulation to scalp EEG changes: a biophysical and physiological modeling study. PLoS One, 8(2):e57330, February 2013.

[10] G Ruffini, R Sanchez-Todo, L Dubreuil, R Salvador, D Pinotsis, E K Miller, F Wendling, E. Santarnecchi, and A Bastos. P118 a biophysically realistic laminar neural mass modeling framework for transcranial current stimulation. Clin. Neurophysiol., 131(4):e78–e79, April 2020.

[11] Roser Sanchez-Todo, André M Bastos, Borja Mercadal, Edmundo Lopez Sola, Maria Guasch, Fabrice Wendling, Emiliano Santarnecchi, and Giulio Ruffini. A physical neural mass modeling framework for laminar cortical circuits in brain stimulation. Brain Stimul., 14(6):1592, November 2021.

[12] Edmundo Lopez-Sola, Roser Sanchez-Todo, Elia Lleal, Elif Köksal-Ersöz, Maxime Yochum, Julia Makhalova, Borja Mercadal, Ricardo Salvador, Diego Lozano-Soldevilla, Julien Modolo, Fabrice Bartolomei, Fabrice Wendling, Pascal Benquet, and Giulio Ruffini. A personalizable autonomous neural mass model of epileptic seizures. bioRxiv, December 2021.

[13] Paul L Nunez and Ramesh Srinivasan. Electric Fields of the Brain: The neurophysics of EEG. Oxford University Press, London, England, 2 edition, 2006.

[14] Henrik Lindén, Espen Hagen, Szymon Leski, Eivind S. Norheim, Klas H. Pettersen, and Gaute T. Einevoll. LFPy: A tool for biophysical simulation of extracellular potentials generated by detailed model neurons. Frontiers in Neuroinformatics, 7(JAN):1–15, 2014.

[15] Espen Hagen, David Dahmen, Maria L. Stavrinou, Henrik Lindén, Tom Tetzlaff, Sacha J. Van Albada, Sonja Grün, Markus Diesmann, and Gaute T. Einevoll. Hybrid scheme for modeling local field potentials from point-neuron networks. Cerebral Cortex, 26(12):4461–4496, 2016.

[16] Alberto Mazzoni, Henrik Lindén, Hermann Cuntz, Anders Lansner, Stefano Panzeri, and Gaute T. Einevoll. Computing the Local Field Potential (LFP) from Integrate-and-Fire Network Models. PLoS Computational Biology, 11(12):1–38, 2015.

[17] Espen Hagen, Solveig Næss, Torbjørn V. Ness, and Gaute T. Einevoll. Multimodal modeling of neural network activity: Computing LFP, ECoG, EEG, and MEG signals with LFPy 2.0. Frontiers in Neuroinformatics, 12(December), 2018.

[18] Pablo Martínez-Cañada, Torbjørn V. Ness, Gaute T. Einevoll, Tommaso Fellin, and Stefano Panzeri. Computation of the electroencephalogram (EEG) from network models of point neurons. PLoS Computational Biology, 17(4):1–41, 2021.

[19] Solveig Næss, Geir Halnes, Espen Hagen, Donald J. Hagler, Anders M. Dale, Gaute T. Einevoll, and Torbjørn V. Ness. Biophysically detailed forward modeling of the neural origin of EEG and MEG signals. NeuroImage, 225(September 2020):117467, 2021.

[20] Wilfrid Rall. Core Conductor Theory and Cable Properties of Neurons. In ER Kandel, JM Brookhart, and VB Montcastle, editors, Comprehensive Physiology, pages 39–97. Wiley, Bethesda, MD, USA, 1977.

[21] Nicholas T. Carnevale and Michael L. Hines. The NEURON Book. Cambridge University Press, 2006.

[22] Julian M.L. Budd, Krisztina Kovács, Alex S. Ferecskó, Péter Buzás, Ulf T. Eysel, and Zoltán F. Kisvarday. Neocortical axon arbors trade-off material and conduction delay conservation. PLoS Computational Biology, 6(3), 2010.

[23] Mainen ZF and Sejnowski TJ. Influence of dendritic structure on firing pattern in model neocortical neurons. Nature, 382(July):363–366, 1996.

[24] Fuyuki Karube, Katalin Sári, and Zoltán F. Kisvárday. Axon topography of layer 6 spiny cells to orientation map in the primary visual cortex of the cat (area 18). Brain Structure and Function, 222(3):1401–1426, 2017.

[25] Andres Carvallo, Julien Modolo, Pascal Benquet, Stanislas Lagarde, Fabrice Bartolomei, and Fabrice Wendling. Biophysical Modeling for Brain Tissue Conductivity Estimation Using SEEG Electrodes. IEEE Transactions on Biomedical Engineering, 66(6):1695–1704, 2019.

[26] Michael A. Moffitt and Cameron C. McIntyre. Model-based analysis of cortical recording with silicon microelectrodes. Clinical Neurophysiology, 116(9):2240–2250, 2005.

[27] Christine Haberler, François Alesch, Peter R. Mazal, Peter Pilz, Kurt Jellinger, Michaela M. Pinter, Johannes A. Hainfellner, and Herbert Budka. No tissue damage by chronic deep brain stimulation in Parkinson’s disease. Annals of Neurology, 48(3):372–376, 2000.

[28] B. H. Jansen and V. G. Rit. Electroencephalogram and visual evoked potential generation in a mathematical model of coupled cortical columns. Biol Cybern, 73(4):357–66, 1995.

[29] Kenneth D. Harris and Thomas D. Mrsic-Flogel. Cortical connectivity and sensory coding. Nature, 503(7474):51–58, 2013.

[30] Sean L. Hill, Yun Wang, Imad Riachi, Felix Schurmann, and Henry Markram. Statistical connectivity provides a sufficient foundation for specific functional connectivity in neocortical neural microcircuits. Proceedings of the National Academy of Sciences of the United States of America, 109(42), 2012.

[31] Shingo Murakami and Yoshio Okada. Invariance in current dipole moment density across brain structures and species: Physiological constraint for neuroimaging. NeuroImage, 111(5):49–58, 2015.

[32] Henry Markram, Eilif Muller, Srikanth Ramaswamy, Michael W. Reimann, Marwan Abdellah, Carlos Aguado Sanchez, Anastasia Ailamaki, Lidia Alonso-Nanclares, Nicolas Antille, Selim Arsever, Guy Antoine Atenekeng Kahou, Thomas K. Berger, Ahmet Bilgili, Nenad Buncic, Athanassia Chalimourda, Giuseppe Chindemi, Jean Denis Courcol, Fabien Delalondre, Vincent Delattre, Shaul Druckmann, Raphael Dumusc, James Dynes, Stefan Eilemann, Eyal Gal, Michael Emiel Gevaert, Jean Pierre Ghobril, Albert Gidon, Joe W. Graham, Anirudh Gupta, Valentin Haenel, Etay Hay, Thomas Heinis, Juan B. Hernando, Michael Hines, Lida Kanari, Daniel Keller, John Kenyon, Georges Khazen, Yihwa Kim, James G. King, Zoltan Kisvarday, Pramod Kumbhar, Sébastien Lasserre, Jean Vincent Le Bé, Bruno R.C. Magalhães, Angel Merchán-Pérez, Julie Meystre, Benjamin Roy Morrice, Jeffrey Muller, Alberto Muñoz-Céspedes, Shruti Muralidhar, Keerthan Muthurasa, Daniel Nachbaur, Taylor H. Newton, Max Nolte, Aleksandr Ovcharenko, Juan Palacios, Luis Pastor, Rodrigo Perin, Rajnish Ranjan, Imad Riachi, José Rodrigo Rodríguez, Juan Luis Riquelme, Christian Rössert, Konstantinos Sfyrakis, Ying Shi, Julian C. Shillcock, Gilad Silberberg, Ricardo Silva, Farhan Tauheed, Martin Telefont, Maria Toledo-Rodriguez, Thomas Tränkler, Werner Van Geit, Jafet Villafranca Díaz, Richard Walker, Yun Wang, Stefano M. Zaninetta, Javier Defelipe, Sean L. Hill, Idan Segev, and Felix Schürmann. Reconstruction and Simulation of Neocortical Microcircuitry. Cell, 163(2):456–492, 2015.

[33] Nicole A. Young, Christine E. Collins, and Jon H. Kaas. Cell and neuron densities in the primary motor cortex of primates. Frontiers in Neural Circuits, 7(FEBRUARY 2013):1–11, 2013.

[34] Idan Segev, John Rinzel, and Gordon M Shepherd. The theoretical foundation of dendritic function. Computational Neuroscience Series. Bradford Books, Cambridge, MA, November 1994.

[35] Christof Koch and Idan Segev, editors. Methods in neuronal modeling. Computational Neuroscience Series. Bradford Books, Cambridge, MA, 2 edition, January 2003.

[36] Guy Eyal, Matthijs B. Verhoog, Guilherme Testa-Silva, Yair Deitcher, Ruth Benavides-Piccione, Javier DeFelipe, Christiaan P. J. de Kock, Huibert D. Mansvelder, and Idan Segev. Human Cortical Pyramidal Neurons: From Spines to Spikes via Models. Frontiers in Cellular Neuroscience, 12(June):1–24, jun 2018.

[37] Delphine Cosandier-Rimélé, Jean Michel Badier, Patrick Chauvel, and Fabrice Wendling. A physiologically plausible spatio-temporal model for EEG signals recorded with intracerebral electrodes in human partial epilepsy. IEEE Transactions on Biomedical Engineering, 54(3):380–388, 2007.

[38] Cristopher M. Niell and Michael P. Stryker. Highly selective receptive fields in mouse visual cortex. Journal of Neuroscience, 28(30):7520–7536, 2008.

[39] Yuta Senzai, Antonio Fernandez-Ruiz, and György Buzsáki. Layer-Specific Physiological Features and Interlaminar Interactions in the Primary Visual Cortex of the Mouse. Neuron, 101(3):500-513.e5, 2019.

[40] Xiaolong Jiang, Shan Shen, Cathryn R. Cadwell, Philipp Berens, Fabian Sinz, Alexander S. Ecker, Saumil Patel, and Andreas S. Tolias. Principles of connectivity among morphologically defined cell types in adult neocortex. Science, 350(6264), 2015.

[41] Robin Tremblay, Soohyun Lee, and Bernardo Rudy. GABAergic Interneurons in the Neocortex: From Cellular Properties to Circuits. Neuron, 91(2):260–292, 2016.

